# Emergence of a novel *Salmonella enterica* serotype Reading clone is linked to its expansion in commercial turkey production, resulting in unanticipated human illness in North America

**DOI:** 10.1101/855734

**Authors:** Elizabeth A. Miller, Ehud Elnekave, Cristian Flores Figueroa, Abigail Johnson, Ashley Kearney, Jeannette Munoz Aguayo, Kaitlin Tagg, Lorelee Tschetter, Bonnie Weber, Celine Nadon, Dave Boxrud, Randall S. Singer, Jason P. Folster, Timothy J. Johnson

## Abstract

Concurrent separate human outbreaks of *Salmonella enterica* serotype Reading occurred in 2017-2019 in the United States and Canada, which were both linked to the consumption of raw turkey products. In this study, a comprehensive genomic investigation was conducted to reconstruct the evolutionary history of *S.* Reading from turkeys, and to determine the genomic context of outbreaks involving this rarely isolated *Salmonella* serotype. A total of 988 isolates of U.S. origin were examined using whole genome-based approaches, including current and historical isolates from humans, meat, and live food animals. Broadly, isolates clustered into three major clades, with one apparently highly adapted turkey clade. Within the turkey clade isolates clustered into three subclades, including an “emergent” clade that only contained isolates dated 2016 or later, including many of the isolates from these outbreaks. Genomic differences were identified between emergent and other turkey subclades suggesting that the apparent success of currently circulating subclades clade is, in part, attributable to plasmid acquisitions conferring antimicrobial resistance, gain of phage-like sequences with cargo virulence factors, and mutations in systems that may be involved in beta-glucuronidase activity and resistance towards colicins. U.S. and Canadian outbreak isolates were found interspersed throughout the emergent subclade and the other circulating subclade. The emergence of a novel *S*. Reading turkey subclade, coinciding temporally with expansion in commercial turkey production and with U.S. and Canadian human outbreaks, indicates that emergent strains with higher potential for niche success were likely vertically transferred and rapidly disseminated from a common source.

**Importance:** Increasingly, outbreak investigations involving foodborne pathogens are confounded by the inter-connectedness of food animal production and distribution, necessitating high-resolution genomic investigations to determine their basis. Fortunately, surveillance and whole genome sequencing, combined with the public availability of these data, enable comprehensive queries to determine underlying causes of such outbreaks. Utilizing this pipeline, it was determined that a novel clone of *Salmonella* Reading has emerged that coincides with increased abundance in raw turkey products and two outbreaks of human illness in North America. The rapid dissemination of this highly adapted and conserved clone indicates that it was likely obtained from a common source and rapidly disseminated across turkey production. Key genomic changes may have contributed to its apparent continued success in the barn environment, and ability to cause illness in humans.

## Introduction

*Salmonella enterica* subsp. *enterica* derived from poultry meat continue to serve as a primary cause of salmonellosis infections towards humans in the United States (1, 2). Among the more than 2500 serotypes that have been identified to date, only a handful of them consistently top the list as those causing the majority of cases of human illness. Estimates on human salmonellosis cases from poultry in the U.S. vary depending on the method used from 10-29.1%, and specifically from turkeys 5.5% (3, 4).

*S*. Reading is a serotype of *S. enterica* subsp. *enterica* first identified in 1916 from a water supply in Reading, England (5), and subsequently identified in various animal hosts, including poultry (6–10). Human outbreaks due to *S*. Reading historically have been rare. In 1956-1957, an outbreak involving *S*. Reading occurred in the U.S., sickening 325 people across multiple states (11). In 2008, 30 persons were involved in an outbreak linked to iceberg lettuce in Finland (12). In 2014-2015, an outbreak of unknown origin was described, with 31 confirmed cases in Canada involving persons of Mediterranean descent (13).

Commercial turkey production is commonly identified as a primary reservoir of *S*. Reading (2, 14–17). Given its low isolation frequency, relatively little is known about the biology of *S*. Reading compared with other serotypes. With that said, *S*. Reading has been shown to have enhanced ability to form biofilms under stress conditions (18) and seems to have the ability to withstand environmental conditions, as they have been isolated from produce (19). Multidrug resistance phenotypes, including resistance towards third-generation cephalosporins, also appear to be common in *S*. Reading strains, including those in dairy cows and beef feedlot cattle (20–22).

Two separate, large outbreaks of *S*. Reading were recently reported in North America. In the U.S., the Centers for Disease Control and Prevention declared an outbreak from November 2017 through March 2019 (23), although human cases of salmonellosis due to *S*. Reading have continued to date. The outbreak was linked to live turkeys and raw turkey products, but no single source product or company was attributed to the entire outbreak. This outbreak resulted in 358 illnesses, 133 hospitalizations, and 1 death across 43 states. In Canada, a separate multi-province outbreak was declared in October 2018 by the Public Health Agency of Canada and is currently active (24). To date (October 2019), there have been 110 identified cases.

Given the widespread nature of these recent North American *S*. Reading outbreaks, there is a pressing need to better understand the ecology and evolution of this foodborne pathogen within suspected animal reservoirs. As such, the purpose of this study was to perform a comprehensive genomic investigation to reconstruct the evolutionary history of *S*. Reading, and to determine if underlying genomic changes within *S*. Reading correlated with outbreaks involving this rarely isolated *Salmonella* serotype.

## Results

### *S.* Reading isolates cluster phylogenetically by host source

Using assembled sequences (*n* = 988) from human illness, meat products, live animals, and environmental sources, isolates were first assigned to 7-gene multilocus sequence types (STs) using the scheme from the PubMLST website (https://pubmlst.org) (25). Based on this scheme, six STs were identified with three dominating: one containing primarily turkey-source and human-source isolates (ST412; 83.7% of isolates), one containing primarily swine/bovine-source and human-source isolates (ST1628; 10.2% of isolates), and one containing primarily human-source isolates (ST93; 5.8% of isolates) (Figure 1). Animal host source was strongly correlated with ST, with 99.6% (564/566) of total turkey-source isolates belonging to ST412 and 93.8% (45/48) and 84.1% (37/44) of swine-source and bovine-source isolates belonging to ST1628, respectively. To rule out temporal bias in the clustering of same host-source isolates by ST, isolates were also characterized based on year of isolation using the same ST scheme (Figure S1). This demonstrated evenness with regard to isolation date across the major STs.

**Figure 1.**
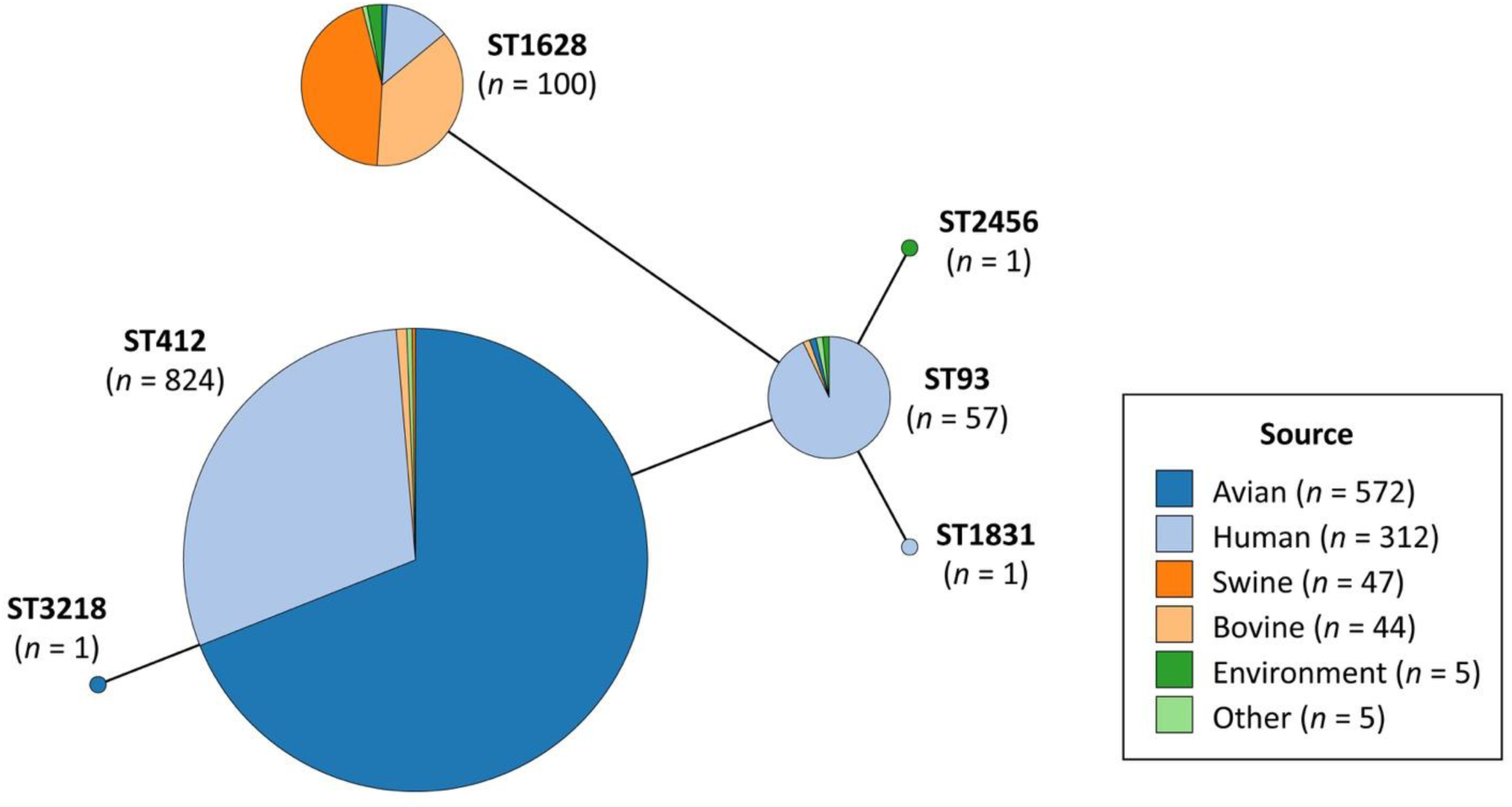
Minimum spanning tree of STs using the Achtman 7-gene MLST scheme for 985 *S*. Reading isolates. Three isolates (swine-, chicken-, and human-source) are not included because their STs could not be determined. Tree is colored based on isolate host source.

Core genome MLST (cgMLST) profiles based upon 3,002 loci were then identified for all isolates, allowing for up to either two allelic differences (Figure S1) or five allelic differences (Figure S1). In all analyses, there was clear and consistent separation based upon animal host source, separating isolates into three major groups.

To gain further resolution, a whole genome SNP-based phylogenetic tree was constructed for all isolates (Figure 2). The resulting tree contained 11086 core SNPs and resolved isolates into three primary clades (designated Clades 1-3), corresponding to MLST and cgMLST results. Clade 1 (*n* = 828) was comprised mainly of turkey-source and human-source isolates, and all but one turkey-source isolate fell within this clade. Clade 2 (*n* = 59) was primarily human-source isolates. Clade 3 (*n* = 101) contained mainly swine-source and bovine-source isolates, with 95.8% (46/48) and 84.1% (37/44) of total swine-source and bovine-source isolates falling within this clade, respectively. Average core SNP distances were investigated between clades (Table S1), revealing that Clades 1-2 were more similar to one another (mean core SNP difference 1638.72 ± 8.49) than Clades 1-3 (8165.04 ± 10.91) or Clades 2-3 (9246.30 ± 12.72). Additionally, mean SNP differences for isolates within Clade 1 (7.72 ± 5.61) were lower than those within Clade 2 (59.23 ± 44.10) or Clade 3 (32.87 ± 16.84). To confirm these results were not due to different sample sizes between clades, average core SNP distances were recalculated on a random subsample of each clade (Table S1).

**Figure 2.**
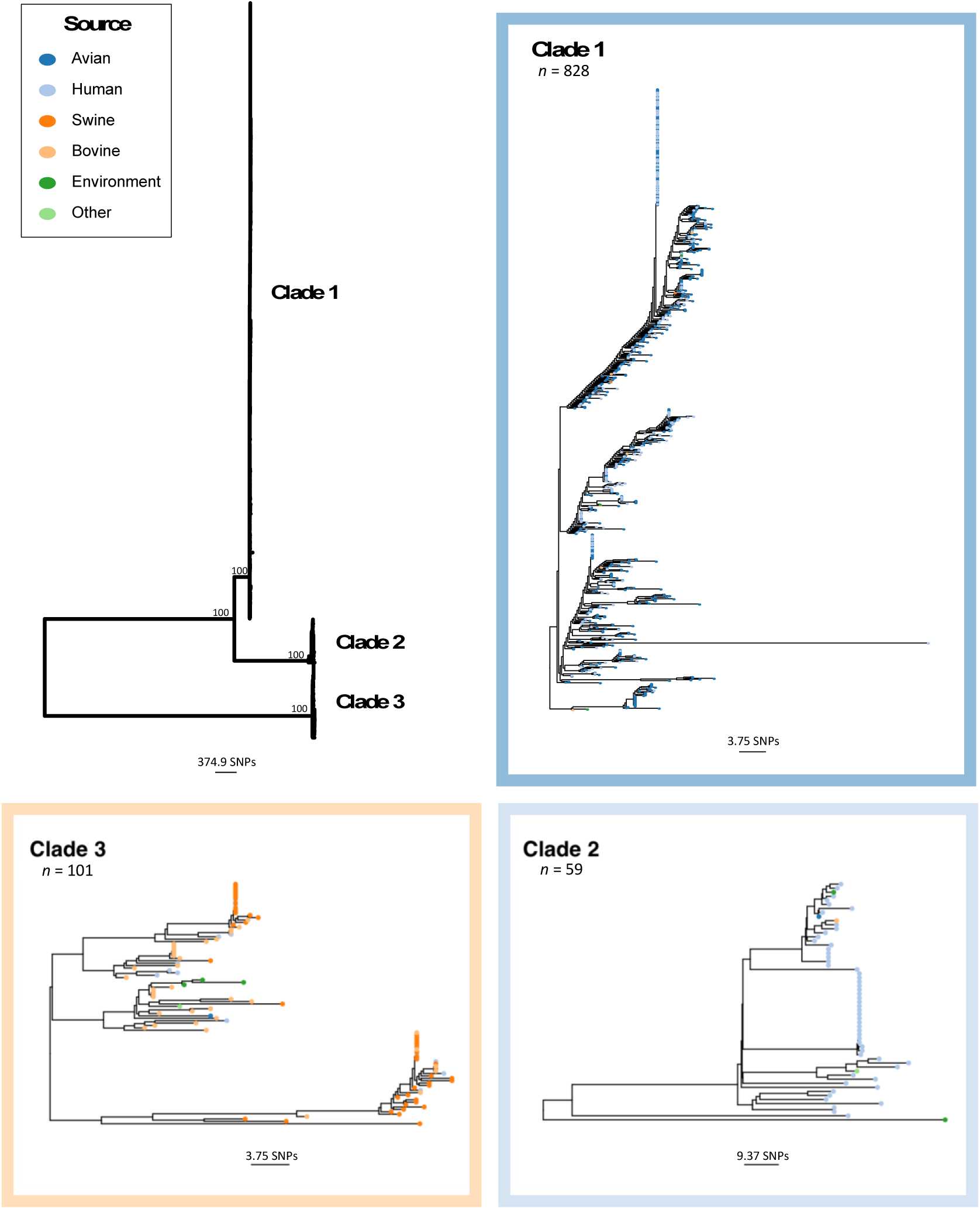
Midpoint-rooted phylogenetic tree of *S*. Reading isolates (*n* = 988) based on core SNPs in nonrecombinant genome regions. All isolates fell into one of three clades: Clade 1 (dark blue; primarily turkey- and human-source), Clade 2 (light blue; primarily human-source), and Clade 3 (orange; primarily swine- and bovine-source). Bootstrap values are shown at the branches differentiating between clades. To allow for more fine-scale view of clade topology, insets show each clade independently (note the difference in scale bars). The color of tip circles indicates isolate host source.

Genome sizes also varied between the three clades, with Clade 2 containing the smallest genomes (median 4.53 ± 0.095 Mb), which were on average 114.50 Kb smaller than Clade 1 genomes (median 4.64 ± 0.050 Mb) and 396.25 Kb smaller than Clade 3 genomes (median 4.92 ± 0.10 Mb) (Figure S2).

A pan-genome approach was then used to investigate specific genomic differences between isolates from Clades 1 and 3, representing the majority of isolates from turkey and swine/bovine sources, respectively. A total of 11366 gene clusters were identified across all 988 isolates, with 3246 (28.6%) present in 100% of isolates (i.e. the “core” genes). Using a cutoff requirement of 100% prevalence vs. 0% prevalence in the two populations, a total of 225 gene clusters were identified as unique to Clade 1, and 180 gene clusters unique to Clade 3 (Supplementary Dataset, https://figshare.com/articles/Supplementary_dataset/10781795). Clade 1 isolates had 15 unique fimbrial system component genes clustered across three systems, including *yadK-L-M-N-V*, *yehA-D,* and a novel K88-like fimbrial system, all of which were inserted in separate genomic locations with genes for each respective system clustered together. Clade 1 isolates also uniquely possessed *prgH-I-K* and *orgA-B*, which are components of the *Salmonella* pathogenicity-associated island SPI-1 (26), genes annotated as cytolethal distending toxin *cdtA-B*, and several prophage-like elements. Conversely, Clade 3 isolates possessed a number of unique fimbrial-like and prophage-like elements compared to those from Clade 1. Also unique to Clade 3 isolates were systems predicted to be involved in type I restriction modification, phosphotransferase activity, and CRISPR/Cas activity.

### A recently emerged clade exists among turkey-source *S.* Reading isolates

The turkey-source isolates from Clade 1 were then examined alone to gain further insight towards their evolution over time. All of these isolates (*n* = 565), except one, belonged to ST412 and were examined at higher resolution using a core SNP-based phylogenetic tree (Figure 3). The core SNP-based phylogenetic tree contained 1093 informative variant sites, and from this, three major clades were designated based upon tree clustering and dates of isolation. The “historical” clade (orange clade in Figure 3, *n* = 65) contained isolates dating 1999-2008. The “contemporary” clade (purple clade in Figure 3, *n* = 201) contained isolates dating 2009-2019, with the majority from 2009-2016. Finally, the “emergent” clade (blue clade in Figure 3, *n* = 295) contained isolates all dating 2017-2019, except for one from 2016. Four isolates were not assigned to a specific clade due to their intermediate location between the contemporary and emergent clades (black clade in Figure 3).

**Figure 3.**
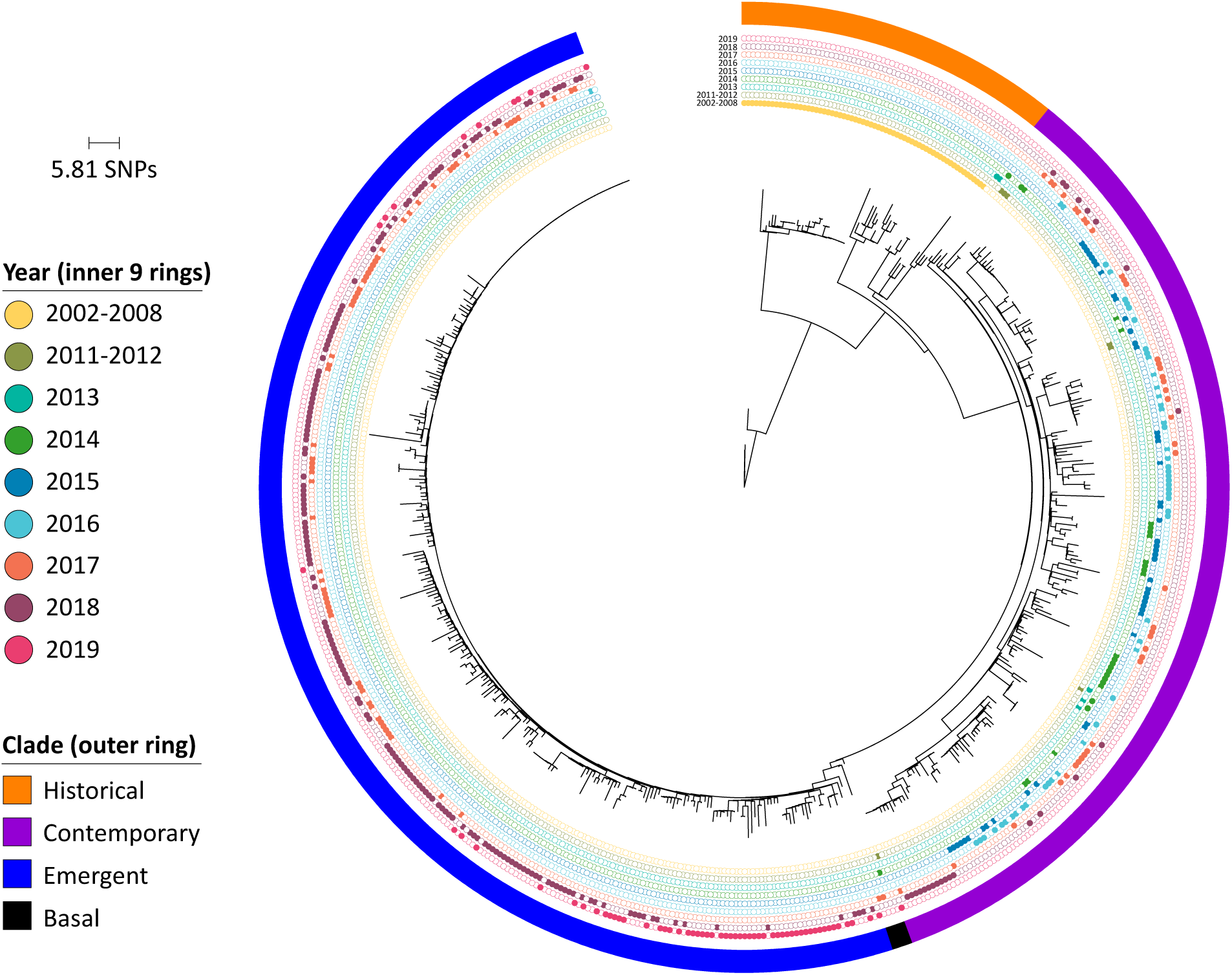
Phylogenetic tree of turkey-source *S*. Reading isolates (*n* = 565) based on core SNPs in nonrecombinant genome regions. The majority of isolates were grouped based on clustering and isolation dates into three clades shown in the outer ring. The inner 9 rings show dates of isolation, with filled circles depicting date for an individual isolate. The tree is rooted with an isolate collected in 2002 (SRR1195634).

The same three-clade structure was also observed in a minimum spanning tree from cgMLST data allowing for up to two allelic differences (Figure 4), where isolates clearly separated by clade designation (historical, contemporary, and emergent) and 57.6% of all isolates in the emergent clade were of the same cgMLST profile. A phylogenetic tree constructed from core genome SNPs and a dendrogram based on hierarchical clustering of all pan-genome genes also showed isolates clustered into the same three clades (Figure S2).

**Figure 4.**
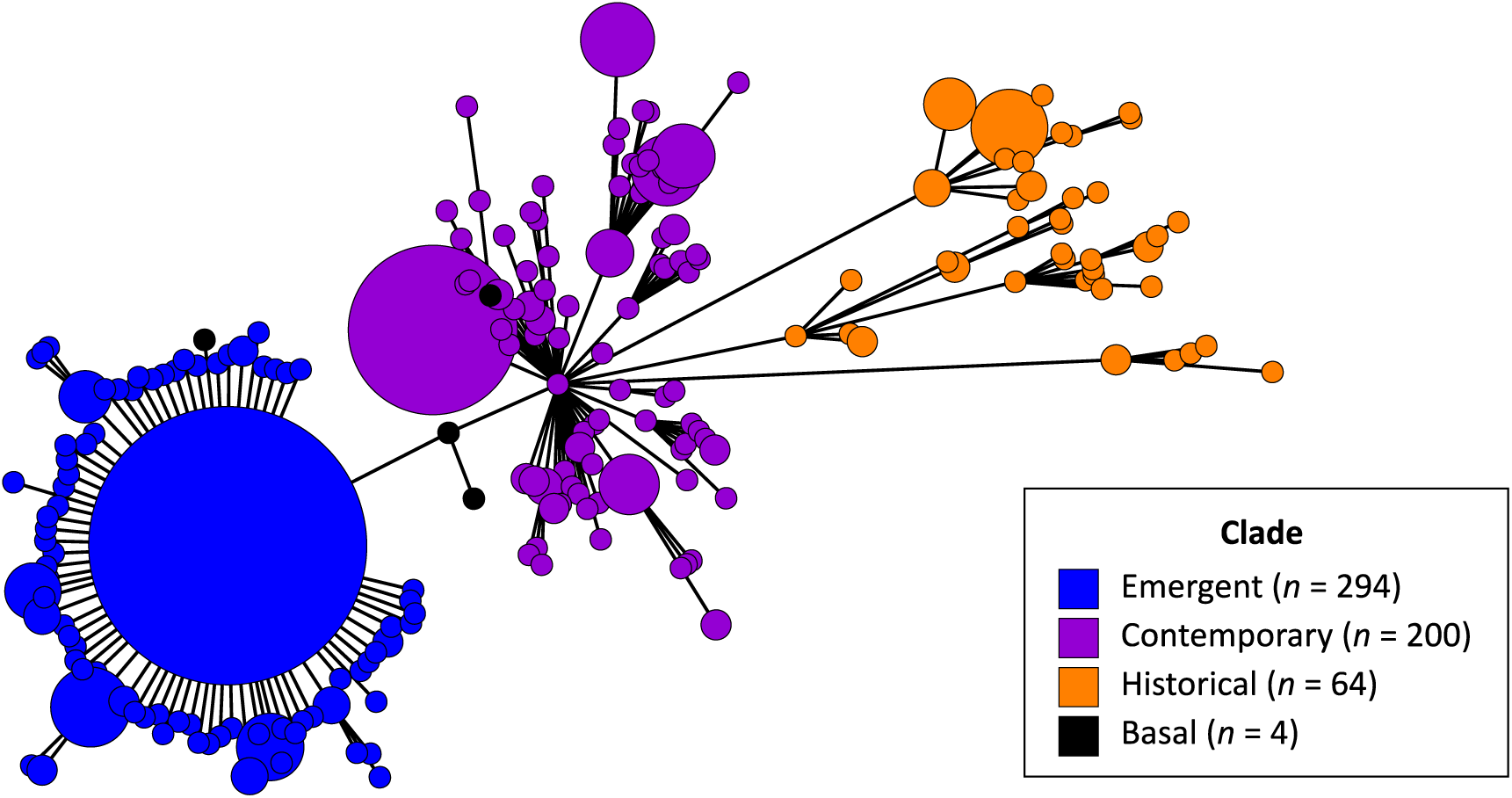
Minimum spanning tree of all turkey-source isolates using the core genome sequence typing (cgMLST) scheme allowing for up to two allelic differences. Three isolates are not included because their cgMLST profiles could not be determined. Tree colors are based on core SNP-based phylogenetic tree clade designations (see Figure 3). Four isolates not assigned to a specific clade are classified as basal to the emergent clade (gray color).

Based upon average core SNP distances (Table 1), the emergent and contemporary clades were more similar to each other (mean core SNP difference 14.35 ± 3.08) than emergent vs. historical (39.95 ± 11.38) or contemporary vs. historical (42.58 ± 11.59). Within clades, emergent clade isolates were more similar to each other (4.67 ± 2.13) than were isolates from the contemporary (10.92 ± 3.88) or historical clades (33.63 ± 18.37).

**Table 1.** Comparison of mean core SNP differences between unique core SNP profiles in the same and different turkey-only phylogenetic clades.

| Clade Comparison | Mean SNP Difference<br>± Std. Dev. | Minimum SNP<br>Difference | Maximum SNP<br>Difference |
| --- | --- | --- | --- |
| All profiles (Historical clade, $n = 44$; Contemporary clade, $n = 151$; Emergent clade, $n = 200$) | | | |
| Overall | $16.74 \pm 14.04$ | 1 | 78 |
| Emergent | $4.67 \pm 2.13$ | 1 | 18 |
| Contemporary | $10.92 \pm 3.88$ | 1 | 23 |
| Historical | $33.63 \pm 18.37$ | 1 | 73 |
| Emergent vs. Contemporary | $14.35 \pm 3.08$ | 6 | 29 |
| Emergent vs. Historical | $39.95 \pm 11.38$ | 23 | 78 |
| Contemporary vs. Historical | $42.58 \pm 11.59$ | 21 | 77 |
| Random profile subset ($n = 44$ samples per clade) | | | |
| Overall | $27.18 \pm 17.51$ | 1 | 76 |
| Emergent | $5.27 \pm 1.95$ | 1 | 12 |
| Contemporary | $10.89 \pm 4.13$ | 1 | 22 |
| Historical | $33.63 \pm 18.37$ | 1 | 73 |
| Emergent vs. Contemporary | $14.61 \pm 3.19$ | 8 | 24 |
| Emergent vs. Historical | $40.18 \pm 11.41$ | 24 | 74 |
| Contemporary vs. Historical | $41.61 \pm 11.60$ | 22 | 76 |

### Small plasmids and associated resistance genes define differences between turkey-source clades

All Clade 1 turkey-source isolates were examined for their possession of genes and mutations known to confer antimicrobial resistance, and plasmid replicons, known among Gram-negative bacteria (Figure 5). When overlaid on the SNP-based phylogenetic tree, several patterns emerged. First, nearly all isolates contained a T57S mutation in *parC* and ColpVC plasmid replicon. An IncQ1 plasmid replicon was found in 20% (41/201) and 33% (98/295) of isolates belonging to the contemporary and emergent clades, respectively. The possession of this plasmid replicon was significantly associated with possession of *sul2*, *tet(A)*, *strA* (*aph(3”)-Ib*), and *strB* (*aph*(*6*)*-Id*) genes conferring the classical SSuT phenotype (Table S2; all pairwise Fisher’s exact test BH-adjusted *P*-values < 0.05). Possession of these traits were found throughout the emergent clade, with some evidence of trait loss scattered infrequently. In contrast, isolates possessing these traits in the contemporary clade were found clustered in the one half of the clade, and were absent from the other half.

**Figure 5.**
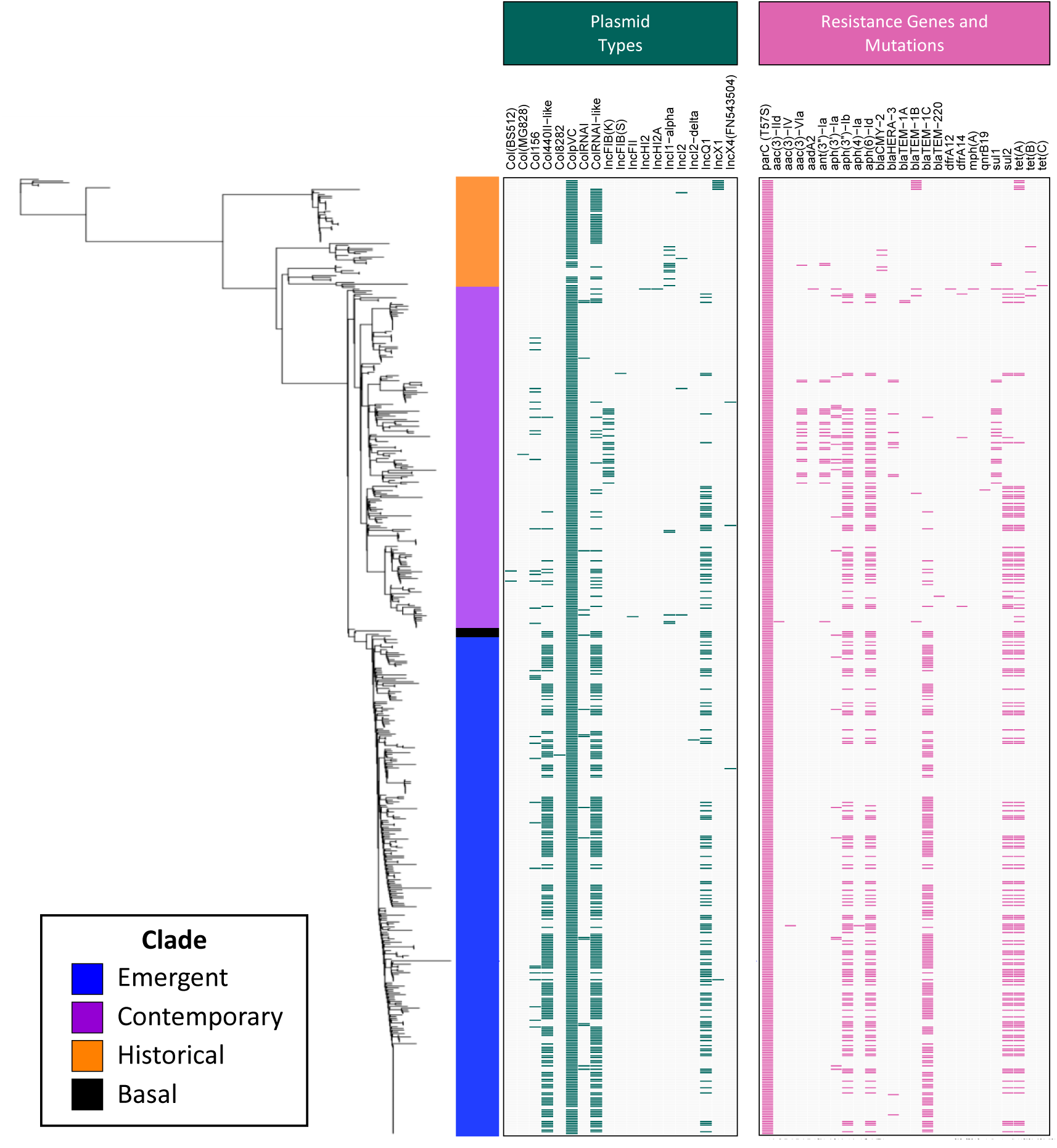
Heatmap displaying the presence of plasmid replicons (dark green) and genes and mutations conferring antimicrobial resistance (pink) across Clade 1 turkey-source isolates.

Isolates belonging to the emergent clade also frequently possessed co-occurring ColRNAI-like and Col440II-like plasmid replicons, present in 61% (181/295) and 65% (191/295) of isolates within this clade. Isolates in the emergent clade were over 20 times more likely to possess both replicons compared to isolates in the historical and contemporary clades (Fisher’s exact test: odds ratio = 0.022, *P*-value < 0.05). Possession of these replicons was also significantly associated with possession of the beta-lactam resistance gene, *bla*_TEM-1C_ (Table S2; all pairwise Fisher’s exact test BH-adjusted *P*-values < 0.05).

Complete sequences of these highly conserved plasmids belonging to IncQ1 and Col440II/ColRNAI-like replicon types were identified and annotated from a representative turkey-source isolate (Figure 6). The IncQ1 replicon and *sul2-strAB-tetAR* genes were co-localized within a 10867-bp mobilizable plasmid containing *mobAC*. The Col440II- and ColRNAI-like replicons were found on a 10384-bp mobilizable plasmid containing *mobAD* and *bla*_TEM-1C_ adjacent to a Tn*2* transposon.

**Figure 6.**
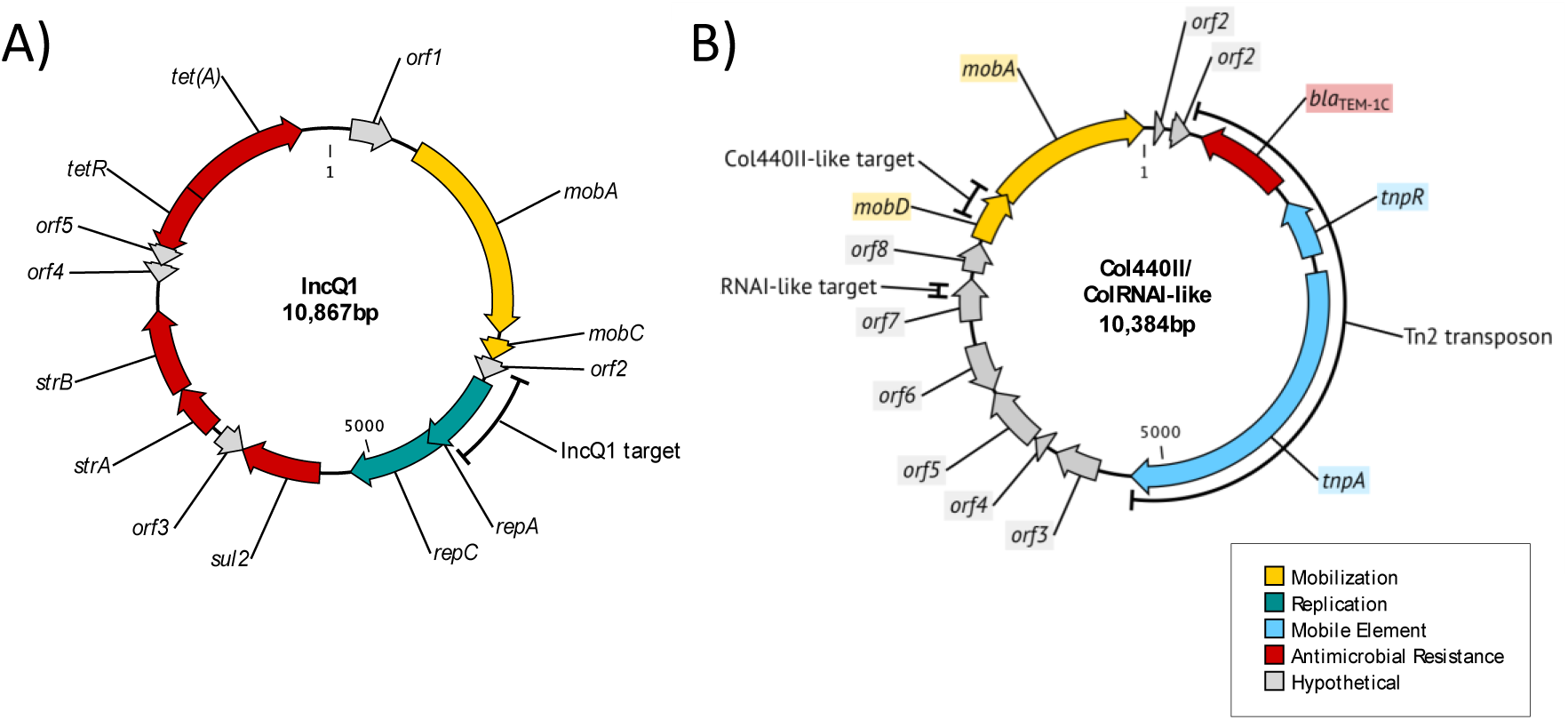
Circular genetic maps of plasmids A) IncQ1 and B) Col440II/ColRNAI-like. Arrows indicate predicted genes and the direction of transcription and are colored to indicate predicted functional category.

To better understand the emergence of this Col440II/ColRNAI plasmid variant, all surveillance data from the European Nucleotide Archive and NCBI SRA databases (until December 2016) was searched for a 282-bp region of the Col440II-like replicon (Figure S3). The first available sequence of the replicon was identified in *S. enterica* in 2000, but did not carry *bla*_TEM-1C_. The first detection of this plasmid replicon carrying *bla*_TEM-1C_ was from a turkey-source *S*. Hadar isolate in 2007, and appearance of this plasmid replicon in *S*. Hadar coincided with subsequent foodborne outbreaks implicating live poultry or poultry products (27, 28), with isolates from those outbreaks containing highly similar plasmids (nucleotide blast of draft assemblies, data not shown). The first detection of this plasmid replicon, including *bla*_TEM-1C_, in *S*. Reading was from a turkey-source isolate in 2014.

### Pangenome-wide association analysis suggests clusters of bacteriophage-associated genes and other elements were gained and lost over time

Comparison of average genome sizes between clades showed an increase in size from the historical clade (median 4.58 ± 0.051 Mb) to the contemporary clade (median 4.66 ± 0.046 Mb) and a subsequent decrease in size to the emergent clade (median 4.63 ± 0.016 Mb) (Figure S2). A pan-genome analysis was used to identify specific genes contributing to this shift in genome size between clades. A total of 6747 gene clusters were produced, of which 4022 (55.8%) were core genes. Of the 2984 accessory genes, the majority (79%) were found in less than 15% of isolates (Figure S3).

Pan-genome-wide association analysis identified 134 genes with significantly differential prevalence between the historical, contemporary, and emergent clades (Figure 7 and Supplementary Dataset, https://figshare.com/articles/Supplementary_dataset/10781795). A large collection of genes primarily encoding bacteriophage-related proteins was absent from the majority of both historical and emergent isolates (< 2.5%), but found in most contemporary isolates (93%) (phage region A in Figure 7). Based on annotations of the representative genome assembly, SRR2407706, all of these genes were clustered in a single region of the *S*. Reading genome (Figure 8; Figure S4), and the majority were homologous to genes from bacteriophages HP1 and HP2. Two separate collections of bacteriophage-related genes were absent from all historical clade isolates, but present in more than 99% of contemporary and emergent clade isolates (phage regions B and C in Figure 7). Both gene clusters could be mapped to separate regions of the *S*. Reading genome (Figure 8; Figure S4), with phage region B genes homologous to genes primarily found in lambda phages GIFSY-1 and GIFSY-2 and phage region C genes homologous to a range of Enterobacteria-specific phages. Of particular note, phage region B included the bacterial virulence-associated gene *sopE* encoding for a type III secretion protein effector, which was surrounded by genes encoding for phage tail and fiber proteins and an IS*L3* family transposase.

**Figure 7.**
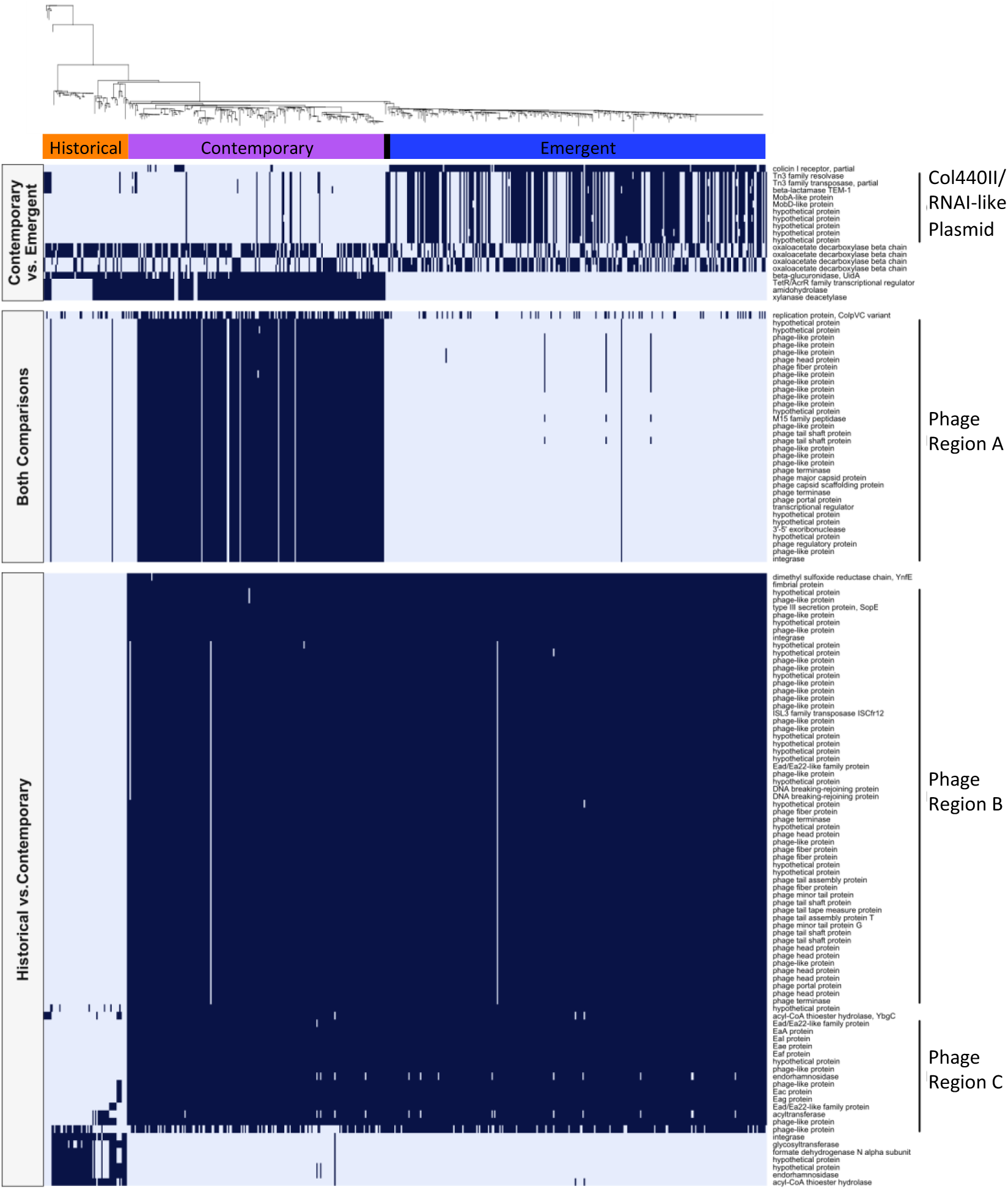
Heatmap displaying the presence (dark blue) and absence (light blue) of genes with significant associations to the historical, contemporary, and/or emergent clades. Left-hand side labels group genes based on the comparison they were identified in: historical vs. contemporary, contemporary vs. emergent, or both comparisons. Right-hand side labels denote genes that clustered into a single region of the *S*. Reading genome.

**Figure 8.**
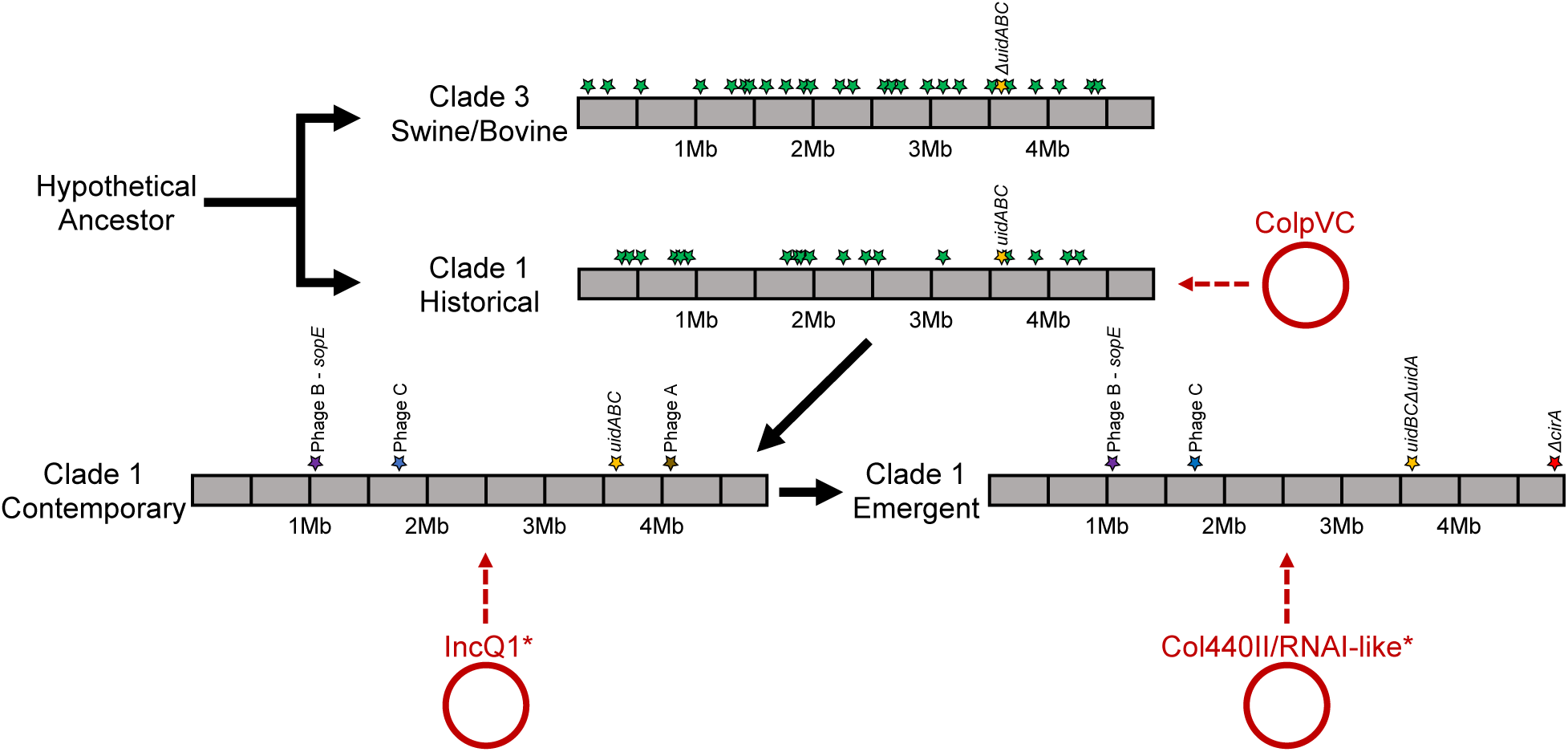
Genetic changes leading from *S*. Reading hypothetical ancestor through the current emergent turkey-source isolate. Green stars indicate unique genomic islands differing between Clades 1 and 3. Purple, blue, and brown stars indicate insertions within Clade 1 contemporary and emergent isolates relative to historical isolates. The gold star indicates insertion of *uidABC*-like region in Clade 1 isolates, where *uidA*-like gene was subsequently truncated in emergent isolates. The red star indicates a truncation of the *cirA* gene in Clade 1 emergent isolates. Plasmid acquisitions are denoted by circles and dashed arrows. *Note that IncQ1 and Col440II/RNAI-like plasmids are found in some other clades, but become dominant in the denoted clades.

Of the 13 genes significantly associated with the emergent clade, 10 were identified as part of the Col440II/RNAI-like plasmid. This included genes encoding TEM-1C beta-lactamase, Tn*3* family transposase and resolvase, mobilization proteins A and D, and five hypothetical proteins. The Col440II-like replicon was significantly more common in isolates from the emergent clade compared to either historical or contemporary clades (all pairwise Fisher’s exact test BH-adjusted *P*-values < 0.05) (Supplementary Dataset, https://figshare.com/articles/Supplementary_dataset/10781795). Additionally, *cirA*, which encodes for a colicin Ia/b receptor, was identified intact in 93.5% of isolates in the contemporary and historical clades, but it was disrupted in the majority (96.9%) of isolates from the emergent clade due to a frameshift insertion of cytosine at position 680. In some of the contemporary clade isolates, amino acids 47-69 of *cirA* were truncated, representing a distinct disruption of CirA compared to the emergent isolates. Similarly, a full-length *uidA*-like gene, which is predicted to encode for a beta-glucuronidase enzyme, was present in 89.8% of contemporary and historical clade isolates, but truncated in all emergent clade isolates. Interestingly, *uidABC* was also found to be absent from Clade 3 isolates in unique fashion compared to Clade 1 emergent isolates.

### Time-scaled phylogenetic analysis

A time-scaled phylogeny of turkey-source sequences (*n* = 398 after removal of duplicated sequences) was reconstructed using a GTR nucleotide substitution model, an uncorrelated lognormal relaxed molecular clock, and a constant growth coalescent model (Figure S4). The model predicted an evolutionary rate of 4.14×10^-7^ substitutions/site/year (higher posterior density (HPD_95%_) = 3.60-4.77×10^-7^) and time to most recent common ancestor (TMRCA) for the Turkey clade was dated to 1984 (1975-1992). The branching of the contemporary and emergent clades was dated to 1997 (1994-1997) with the emergent clade arising in 2015 (2014-2016).

### North American *S.* Reading outbreak isolates cluster with both contemporary and emergent clade turkey-source isolates

To investigate the two recent North American *S*. Reading outbreaks in the context of turkey-source *S*. Reading strains, a core SNP-based phylogenetic tree was constructed for all Clade 1 turkey-source isolates (*n* = 565) and human-source isolates identified as part of the 2017-19 *S*. Reading outbreaks in the U.S. (*n* = 139) and Canada (*n* = 111) (Figure S5). Outbreak isolates from both countries were found clustered with turkey-source isolates from both the contemporary and emergent clades. Specifically, for the U.S. outbreak isolates, 29.5% (41/139) of isolates clustered with the contemporary clade and 69.1% (96/139) with the emergent clade. For Canadian outbreak isolates, the distribution was more balanced between clades, with 52.3% (53/111) clustering with the contemporary clade and 47.7% (58/111) with the emergent clade. A subset of both U.S. and Canadian outbreak isolates shared identical core SNP profiles with some turkey-source isolates. In particular, one prevalent identical SNP profile was found in 96 isolates, including 56 turkey-source isolates, 28 U.S. outbreak isolates, and 12 Canadian outbreak isolates. Mining of CDC and Minnesota of Department of Health data suggests an increase in *S*. Reading starting in 2014 involving Clade 1 contemporary isolates, which was exacerbated by Clade 1 emergent isolates starting in 2016 (Table 2).

**Table 2.** Human cases of *S*. Reading reported by CDC and Minnesota Department of Health (MNDH), compared with percent of human-source isolates used in this study that cluster with turkey-source contemporary or emergent clade isolates. ND = no human case data available.

| Dataset | 2008 | 2009 | 2010 | 2011 | 2012 | 2013 | 2014 | 2015 | 2016 | 2017 | 2018 |
| --- | --- | --- | --- | --- | --- | --- | --- | --- | --- | --- | --- |
| CDC cases | 46 | 53 | 33 | 42 | 58 | 55 | 104 | 139 | 221 | ND | ND |
| MNDH cases | 3 | 3 | 1 | 1 | 3 | 2 | 4 | 7 | 7 | 13 | 21 |
| Cases associated with contemporary clade (%) | ND | ND | ND | ND | 0% | 0% | 82% | 88% | 59% | 44% | 23% |
| Cases associated with emergent clade (%) | ND | ND | ND | ND | 0% | 0% | 0% | 0% | 2% | 29% | 63% |

## Discussion

Multiple outbreaks of *S*. Reading in North America prompted an investigation of the microevolution of this serotype, as human-associated outbreaks due to *S*. Reading are rarely reported. Very clear host separation has occurred between avian-source and bovine/swine-source *S*. Reading isolates, accompanied by large whole genome SNP differences and numerous genomic island differences. This clear separation without intermediate isolates between the two clades (1 vs. 3) suggests that current clades represent host-adapted lineages evolved towards their respective successes in avian vs. bovine/swine hosts. Within the avian clade, a time-scaled phylogeny reconstruction demonstrated the diversification of subclade branches with estimated node ages that align with the current North American outbreaks. In addition, these analyses estimated an evolutionary rate of 4.14×10^-7^ substitutions/site/year, which corresponds to a change of two SNPs per year. The constant population growth selected here may reflect the early stage of this clade’s spread. The data indicate two distinct expansions of *S*. Reading. First, the contemporary clade began the expansion in 2014 with an increased number of human cases compared to previous years (Table 2). In 2017, numbers of human cases again expanded with the surfacing of the emergent clade, coinciding with multiple outbreaks declared in the U.S. and Canada.

Genetic diversity within Clade 1 was lowest compared to all clades studied here, and genetic diversity within the emergent Clade 1 isolates was extremely low. This, combined with dates of isolation, points to the recent emergence of a new clone of *S*. Reading, which was estimated to emerge in 2015 (HPD_95%_:2014-2016) based on the time-scaled phylogeny reconstruction. This emergence coincides with large outbreaks in North America linked to contaminated turkey products, prompting the question of why this strain and serotype has become successful. The overall genetic differences between the avian subclades were subtle, yet may provide important clues highlighting the success of the contemporary and emergent strains. One distinguishing feature of the emergent strains, and contemporary subsets of the circulating clade, was the presence of mobilizable IncQ1 and Col440II/ColRNAI-like small plasmids. Collectively, these plasmids encode resistance towards ampicillin, streptomycin, sulfamethoxazole, and tetracycline. IncQ1 plasmids are broad host range, highly mobilizable plasmids capable of residing in a variety of Gram-negative bacterial species (29). Similar conformations of this plasmid conferring the same SSuT resistance profile have been identified in *S*. Typhimurium in Italy (30). While the presence of these two plasmids appears to be a marker of evolution of the clade, they apparently have been frequently lost by isolates in the emerging clade. There was no association between isolate host source and apparent plasmid loss (i.e., human-versus turkey-source isolates), indicating that plasmid loss is not a function of selective pressure in a particular environment, but instead a function of genetic gain followed by plasmid instability or dispensability.

There was an overall genome size gain between the historical to contemporary/emergent isolates within Clade 1. This was primarily due to acquisition of several phage-like elements within the chromosome. Acquisition of a lambda-like prophage-like element was accompanied by accessory carriage of *sopE* into the contemporary and emergent clades (Figure 8). All avian strains carried the canonical version of *Salmonella* pathogenicity-associated island, SPI-1. SopE, along with SopE2, are guanine nucleotide exchange effector molecules for the type III secretion system encoded by SPI-1 (31). Together, these two molecules are able to act differentially on the RhoGTPase signaling cascade and may promote enhanced inflammatory function. SopE has also been shown to enhance murine colitis (32). SopE has previously been identified on a P2 family phage-like element in *S*. Typhimurium (33) and was associated with persistent epidemic strains in humans and animals. SopE has also been shown to reside on diverse phage types, including lambda-like phage in *S*. Gallinarum, Enteritidis, Hadar and Dublin (34), and was more common in the most common human serotypes in England (35). Therefore, the acquisition of SopE by contemporary and emergent avian clade isolates may represent an advantage for their persistence and virulence.

Two gene disruptions were notable between the emergent and contemporary isolates of Clade 1. First, emergent isolates possessed a frameshift insertion of cytosine at position 680 in the *cirA* gene resulting in a predicted frameshift that was uniform across emergent isolates. Additionally, a portion of the contemporary isolates possessed a truncation of *cirA* that was independent of the mutation identified in emergent isolates. CirA is a catecholate siderophore receptor that also serves as the receptor for colicin ColIb, a pore-forming toxin produced by some *E. coli* and *Salmonella* as a competitive exclusion mechanism (36). ColIb production has been shown to favor producers during competition with ColIb-sensitive strains lacking the plasmid that encodes this system (37). However, mutations in *cirA* have rendered ColIb-sensitive strains resistant to the killing effects of ColIb (38). Furthermore, ColIb is commonly found to reside on IncI1 plasmids, which are ubiquitous among Enterbacteriaceae of commercial turkeys (39, 40). Therefore, it is plausible that disruption of *cirA* in emergent isolates provides a competitive advantage in the gastrointestinal tract against competing ColIb-positive bacteria. Because disruption of this gene was observed convergently in the contemporary and emergent clades, it warrants further study.

A second gene disruption identified among emergent isolates, not present in contemporary or historical isolates, was a deletion of a *uidA*-like sequence accompanied by deletion of an adjacent gene predicted to encode for peptidoglycan deacetylase, PgdA. This region was intact in contemporary and historical isolates. Interestingly, Clade 3 isolates were missing the entire *uidABC* region, but retained *pgdA*. The presence of *uidABC* was sought amongst other phylogenetically proximal *Salmonella* serotypes (41), and was universally present, agreeing with previous studies identifying *Salmonella* clade-specific beta-glucuronidase activity (42). Together, this indicates that the *uidABC* system was ancestrally intact and subsequently truncated/deleted independently in Clade 1 emergent and Clade 3 isolates. The *uidABC* operon encodes enzymes capable of breaking down glucuronidated ligands, freeing them up as a bacterial nutrient source (43). This is typically viewed as a competitive advantage for gut bacteria. However, because these systems were convergently inactivated in two distinct host-adapted clades of *S*. Reading, and beta-glucuronidase systems are known to have a diverse array of functional effects in the gut (44), the possible role of inactivation of this system as a fitness benefit deserves further study.

This study was prompted by two large outbreaks of *S*. Reading in North America linked to the consumption of raw turkey products (23, 24). Our analyses indicate that these outbreaks coincide with the emergence of a novel successful clone of *S*. Reading in North America, and dramatically increased rates of isolation of *S*. Reading in commercial turkey production, independent of company or geographical region. Given these facts, it is quite likely that the introduction of this clone occurred in commercial turkey production rapidly and uniformly. The most parsimonious explanation is that this clone was introduced vertically from a common source. Interestingly, the emergence of this clone coincides with an outbreak of highly pathogenic avian influenza in 2015 that decimated turkey breeder supplies in the upper Midwestern U.S. (45). Thus, the emergence of this clone, combined with rapid repopulation efforts in the turkey industry, may have further contributed to its rapid spread. The microevolution of *S*. Reading in turkeys towards the emergent clade has apparently provided it with evolutionary advantages for success in the growing turkey, the turkey barn environment, and/or the human host. While it is impossible at this time to pinpoint the precise source, this study highlights the power and utility of high-resolution genomics for better understanding the ecology and evolution of outbreaks of foodborne pathogens.

## Materials and methods

### Sample collection and DNA sequencing

Thirty-two isolates from this study were collected from commercial turkey production facilities in the U.S. between October 2016 and October 2018. Samples represent 32 unique premises within multiple turkey-producing companies. Samples were collected by boot sock sampling, environmental swabbing, fluff sampling, or cecal sampling. Enrichments were performed for *Salmonella* by primary enrichment of 1 g sample content in 9 mL in Tetrathionate broth overnight with shaking at 42°C, followed by streaking of the primary enrichment onto XLD agar and incubation overnight at 37°C. Serotyping was performed on isolates following a standard protocol (46). DNA was extracted from cultures using the Qiagen DNeasy kit (Valencia, CS) following manufacturer instructions. Genomic DNA libraries were created using the Nextera XT library preparation kit and Nextera XT index kit v2 (Illumina, San Diego, CA) and sequencing was performed using 2×250-bp dual-index runs on an Illumina MiSeq at the University of Minnesota Mid-Central Research and Outreach Center (Willmar, MN).

### Study population for phylogenomic analysis

A search of NCBI’s Short Read Archive (SRA) was conducted for all available raw sequencing data of isolates annotated as *Salmonella enterica* subsp. *enterica* serotype Reading. Only isolates that met the following criteria were considered: 1) was collected within the United States, 2) had a known isolation year, and 3) had a known isolation source. Raw sequencing reads of all identified isolates (*n* = 989) were downloaded from the SRA using the SRA Toolkit (v2.8.2). An additional 32 isolates collected from U.S. commercial turkey production facilities were sequenced for this study (see *Sample collection and DNA sequencing* for details). A series of quality filtering steps within the bioinformatic processing pipeline (described below) were used to obtain a final sample size of 988 high-quality isolate genomes, including 566 from turkey-related sources (Supplementary Dataset, https://figshare.com/articles/Supplementary_dataset/10781795). A summary of sample filtering steps is depicted in Figure S6.

To investigate the two recent North American *S*. Reading outbreaks in the context of turkey-source *S*. Reading strains, raw sequencing reads from an additional 111 clinical *S*. Reading isolates collected by the Public Health Agency of Canada’s (PHAC) National Microbiology Laboratory were downloaded from the SRA (Supplementary Dataset, https://figshare.com/articles/Supplementary_dataset/10781795). U.S. and Canada clinical isolates were defined as part of the 2017-19 outbreaks based on criteria that included analysis by whole genome sequencing defined by the CDC and PHAC, respectively.

### Genome assembly and quality assessment

All raw FASTQ files were trimmed and quality filtered using Trimmomatic (v0.33) (47), specifying removal of Illumina Nextera adapters, a sliding window of 4 with an average Phred quality score of 20, and 36 as the minimum read length. Trimmed reads were *de novo* assembled using the Shovill pipeline (v1.0.4), which utilizes the SPAdes assembler (48), with default parameters (https://github.com/tseemann/shovill). Assembly quality was assessed with QUAST (v5.0.0) (49). To calculate average sequencing depth of coverage, trimmed reads were mapped to assembled contigs using the BWA-MEM algorithm (v0.7.17) (50) and a histogram of depth was computed using the *genomecov* command in BEDTools (v2.27.1) (51). Only isolates with an N50 ≥ 20000 bps and an average depth of ≥ 20X were included in further analyses (Figure S6).

### Serotype prediction

*In silico* serotype prediction was performed with the *Salmonella In Silico* Typing Resource (SISTR) (v1.0.2) (52). Only isolates with a predicted serotype of Reading for both genoserotyping and cgMLST cluster analysis were included in downstream analyses (Figure S6).

### Sequence typing

*In silico* multilocus sequence typing (MLST) was performed using the software, mlst (v2.16.1) (https://github.com/tseemann/mlst), with the Achtman 7-gene *Salmonella* MLST scheme hosted on the PubMLST website (https://pubmlst.org) (25). Core genome multilocus sequence typing (cgMLST) was performed on the EnteroBase webserver using their custom *Salmonella* cgMLST V2 scheme of 3002 loci (53). Because draft genomes of multiple contigs may frequently contain missing genes, cgMLST profiles were hierarchically clustered allowing for a mismatch of up to two and five alleles. Minimum spanning trees based on both the traditional MLST and cgMLST allelic profiles were generated in Enterobase’s standalone software, GrapeTree (v1.5.0) (54).

### Phylogenetic analysis

Single nucleotide polymorphisms (SNPs) were identified in each sample using Snippy (v4.4.0), with a minimum sequencing depth of 8X (https://github.com/tseemann/snippy) and the *S*. Reading assembly, SRR6374143, as the reference. Separate core SNP alignments were then created for all isolates (*n* = 988) and for all Clade 1 turkey-source isolates (*n* = 565). Based on MLST and cgMLST minimum spanning trees, one turkey isolate clustered separately from all other turkey isolates and was therefore not included in the turkey-source alignment. Recombinant regions were identified with Gubbins (v2.3.4) (55) and masked from the core genome alignments using maskrc-svg (v0.5) (https://github.com/kwongj/maskrc-svg). Samples with >25% missing data were removed from further analyses (Figure S6). The program snp-sites (v2.4.1) was then used to extract all core SNPs and monomorphic sites where the columns did not contain any gaps or ambiguous bases (56). Pairwise core SNP distance matrices were created using snp-dists (v0.6.3) (https://github.com/tseemann/snp-dists) after duplicate core SNP profiles were removed with SeqKit (v0.10.1) (57).

Maximum likelihood trees for both all isolates and the turkey-source isolates only were reconstructed based on the alignments of core SNPs plus monomorphic sites with IQ-TREE (v1.6.10) (58). ModelFinder was used to identify the most appropriate substitution models (59). For the “all-isolate” tree, the model with the best fit according to the Bayesian information criterion was the three substitution-type model (K3Pu) (60) with empirically-derived unequal base frequencies (+F) and the discrete Gamma model of rate heterogeneity model with four rate categories (+G4) (61). For the “turkey-source” tree, the best model was K3Pu+F+I, where the rate heterogeneity model (+I) allowed for a proportion of invariable sites. Branch support for both trees was estimated by performing 1000 ultrafast bootstrap approximation replicates (62). The resulting trees were visualized and annotated using the online tool iTOL (63).

To assess the robustness of clades identified in the turkey-source core SNP-based phylogenetic tree, two additional turkey-source trees were constructed using alternative methods based on the pan-genome (see *Pan-genome analysis* for further details). First, a core genome phylogenetic tree was constructed from the core genome alignment. Core SNPs and monomorphic sites were then extracted from this alignment and used as input into ModelFinder and IQ-TREE. The best model was the transversion substitution model [AG = CT] (TVM) with empirically-derived unequal base frequencies (+F) and allowing for a proportion of invariable sites (+I). Branch support was estimated from 1000 ultrafast bootstrap approximation replicates. Second, a hierarchical clustering dendrogram was generated based on the presence/absence of pan-genome gene clusters. Euclidean distance was calculated using the R package, vegan (v2.5-5) (64), and complete linkage clustering was performed by the *hclust* function from the R package, stats (v3.6.1).

A separate maximum likelihood tree of all Clade 1 turkey-source isolates (*n* = 565) and human-source isolates identified as part of the 2017-19 *S*. Reading outbreaks in the U.S. (*n* = 139) and Canada (*n* = 111) was constructed following the same methods outlined above. As with the turkey-only tree, the best model was identified as K3Pu+F+I, with 1000 ultrafast bootstrap approximation replicates to estimate branch support.

### Time-scaled phylogenetic analysis

Non-duplicate turkey-origin isolates were used. A ‘temporal signal’ of the data was evaluated by generating a linear regression of phylogenetic root-to-tip distances against the sampling dates using Tempest (v1.5) (65), and a positive correlation between root-to-tip distance and collection time (R^2^ = 0.46) was demonstrated. In addition, the ‘temporal signal’ was verified using a tip-date randomization test that was conducted using the package ‘TipDatingBeast’ (v1.0.6) (66) in R (v3.4.3) (67). The evaluated TMRCA for the selected model (below) was compared between the real data and the randomized trials (*n* = 20), and no overlaps were found between the HPD_95%_ intervals and/or mean values (data not shown). A time scaled phylogeny was constructed using BEAST (v 1.10.4) (68). A general time reversible (GTR) substitution model was used for nucleotide substitution and both ‘uncorrelated lognormal relaxed’ and ‘strict’ molecular clocks with different coalescent population models (i.e. constant growth, logistic growth, exponential growth, GMRF Bayesian skyride and Bayesian skyline) were explored, correcting for ascertainment bias. Log marginal likelihoods obtained using path sampling (PS) / stepping-stone sampling (SS) (69, 70) were compared. An evolutionary rate of 2.64 × 10^-7^ mutations per site per year, previously estimated for *S*. I 4,[5],12:i:-ST34 (Elnekave et al, unpublished) was used as the mean estimation for the clock rate prior. Each model combination was tested for at least two independent Markov chain Monte Carlo (MCMC) runs of at least 200 million generations, with sampling every 20000 generations. Convergence and proper mixing of all MCMC runs (effective sample size > 200) and the agreement between two independent MCMC runs of the same model were verified manually in Tracer (v1.7.1) (71) after excluding 10% of the MCMC chain as a burn-in. The model with the highest log Bayes factor value was the GTR-uncorrelated lognormal relaxed-constant population growth combination. LogCombiner (v1.10.4) (68) was used to combine the two independent MCMC runs of the final model after exclusion of 10% burn-in period. Package ggtree (v1.10.5) (72) in R software was used for tree visualization.

### Genetic feature identification

Acquired resistance genes and known chromosomal mutations conferring antibiotic resistance were identified in sample assemblies using staramr (v0.3.0) (https://github.com/phac-nml/staramr) with the ResFinder and PointFinder databases (73, 74). A minimum identity of 90% was used for matching to both databases, with default minimum coverage lengths of 60% for ResFinder and 95% for PointFinder. Plasmid replicon markers were identified using ABRicate (v.0.8.13) (https://github.com/tseemann/abricate) with the PlasmidFinder database (75) and a minimum identity of 90% and minimum coverage length of 60%. ABRicate was also used to screen sample assemblies for the two additional plasmid replicons, Col440II-like and ColRNAI-like (https://github.com/StaPH-B/resistanceDetectionCDC), as they were of interest, but not present in the PlasmidFinder database. A heatmap of the presence and absence of plasmid types and antimicrobial resistance genes was created with the R packages, ggtree (v1.16.4) and tidytree (v0.2.5) (76). To test for significant nonrandom associations between genomic features of interest, one-sided Fisher’s exact tests were performed on 2×2 contingency tables using the R function, *fisher.test*, with the Benjamini-Hochberg (BH) procedure to adjust *P*-values for multiple testing (77).

### Plasmid and accessory element annotation and analysis

Based upon plasmid replicon results, two plasmids were selected belonging to IncQ1 and Col440II/RNAI-like replicons. These completed plasmids were searched via nucleotide BLAST across several isolates within each clade to confirm their conservation. Representative plasmid sequences were used from strain SRR8925563. Genes were predicted using Prokka (78) and plasmids were annotated and visualized via CLC Sequence Viewer (v8.0.0) (Qiagen, Aarhus, Denmark). For clade-to-clade chromosome comparisons, representative genome assemblies were retrieved for Clade 1 - historical (SRR1583085), Clade 1 - contemporary (SRR2407706), Clade 1 - emergent (SRR6904571), and Clade 3 (SRR5865228) and annotated via Prokka. MAUVE (79) was used to re-order chromosomal contigs of the draft assemblies to that of a completed *S*. Reading chromosome (Genbank accession no. CP030214) (80). MAUVE was then used to align representative chromosomes and compare for genomic differences.

### Plasmid prevalence over time and serotypes

To determine the prevalence of the Col440II/ColRNAI-like plasmid in *Salmonella enterica* over time, a 282-bp region of the Col440II-like replicon was used to search the publicly available ENA/SRA databases (through December 2016; 90% threshold) (81). Metadata for sequences positive for the 282-bp target was downloaded from NCBI. Resistance gene content was determined using an in-house database adapted from ResFinder 3.0 (90% identity, 60% cutoff). Sequenced isolates with both serotype and year of collection available were included in the analysis (*n* = 100).

### Pan-genome analyses

Sample assemblies were annotated with Prokka (v1.13.4) and a core genome alignment was generated using Roary (v3.12.0) (82). Coding sequences were clustered into “gene clusters” using the default 95% sequence identity. “Core genes” were defined as gene clusters identified in 100% of isolates, while an “accessory genes” were defined as clusters present in <100% of isolates. A presence/absence matrix heatmap of accessory genes was created using the *roary_plots.py* script (https://github.com/sanger-pathogens/Roary/tree/master/contrib/roary_plots). Scoary (v1.6.16) (83) was then used to conduct a pan-genome-wide association analysis comparing the prevalence of gene clusters between phylogenetic clades. Specifically, in the all-isolate trees, “Clade 1” isolates were compared to “Clade 3” isolates, and in the turkey-source tree, “contemporary clade 1b” isolates were compared separately to both “emergent clade 1c” and “historical clade 1a” isolates. Genes identically distributed across samples were collapsed into a single gene cluster with the *collapse* option. For the turkey-only tree, a gene cluster was reported as significantly associated with a particular clade if it had a BH-adjusted *P*-value ≤ 0.05 and was present in ≥ 60% of isolates in one clade and ≤ 40% in the other clade. The reference sequence(s) of each significant gene cluster were then annotated using the top hit(s) from a BLASTX search against the NCBI’s non-redundant protein sequence database (81). Heatmaps comparing the percent of genomes possessing the significant gene cluster between clades were created using the R package, ggplot2 (v3.2.0) (84). Because not all plasmid replicons of interest were identified by Prokka and thus were not included in the pan-genome analysis, separate 2×2 Fisher’s exact tests were performed for each identified plasmid replicon with BH-adjusted *P*-values. Follow-up annotations of bacteriophage regions in the *S*. Reading genome were conducted on a representative genome assembly from the contemporary clade, SRR2407706, with the web-based phage search tool, PHASTER (85).

## Acknowledgments

The authors would like to thank the turkey producers of the United States for their willingness to collaborate in this study. Isolates and data for outbreak cases in Canada were provided courtesy of the PulseNet Canada Steering Committee and members of the Canadian Public Health Laboratory Network. Bioinformatics were supported using tools available from the Minnesota Supercomputing Institute. Sequencing reagents for this study were donated by the Mid-Central Research and Outreach Center, Willmar, MN, USA.

## Supplementary Figure Legends

**Figure S1.** A) Minimum spanning tree of STs using the Achtman 7-gene MLST scheme for 985 S. Reading isolates. Three isolates (swine-, chicken-, and human-source) are not included because their STs could not be determined. Tree is colored based on isolate collection year. B-C) Minimum spanning tree of STs using the EnteroBase core genome sequence typing (cgMLST) scheme allowing for up to B) two allelic differences and C) five allelic differences. Seven isolates (three turkey-source and four human-source) are not included because their cgMLST profiles could not be determined. Tree is colored based on isolate host source.

**Figure S2.** A-B) Box and whisker plots displaying the distribution of genome sizes in kilobases between A) Clades 1-3 of the all-host phylogenetic tree (*n* = 988) and B) the historical, contemporary, and emergent clades of the turkey-source phylogenetic tree (*n* = 565). The upper and lower edges of the boxes correspond to the first and third quartiles, respectively, and the upper and lower whiskers extend to the largest and smallest values no further than 1.5 * inter-quartile range. Data beyond these whiskers are considered outliers. C-D) Clustering of turkey-source *S*. Reading isolates (*n* = 565) based on C) a phylogenetic tree based on variant sites in the core genome alignment (1072 core SNPs) and D) a complete linkage cluster dendrogram based on Euclidean distances of presence/absence of pan-genome genes. The inner green rings around both trees denote the collection year range of each isolate. The outer rings denote the core SNP-based phylogenetic tree clade designations from Figure 3.

**Figure S3.** A) The presence of the Col440II-like replicon in *Salmonella enterica* by serotype over time. B) Presence-absence heatmap displaying the distribution of accessory genes across turkey-source isolates. Genes were identified by clustering coding sequences based on 95% sequence identity.

**Figure S4.** A-C) Genetic maps of bacteriophage regions of the *S*. Reading genome based on representative genome assembly, SRR2407706. Maps consist primarily of genes identified in the pan-genome-wide association analysis (Figure 7), including A) phage region A, B) phage region B, and C) phage region C. Arrows indicate predicted genes and the direction of transcription and are colored to indicate predicted functional category based on PHASTER annotations. D) A time-scaled phylogeny of turkey sequences (*n* = 398 after removal of duplicated sequences). Tips are annotated according to the main clades and the main nodes ages (HPD_95%_) are indicated.

**Figure S5.** Phylogenetic tree of *S*. Reading isolates from turkey (*n* = 565) and humans (*n* = 250) based on 1242 core SNPs in nonrecombinant genome regions. Only human-source isolates classified as part of the 2017-19 *S*. Reading outbreaks in the U.S. and Canada were included. The inner colored ring around the tree denotes the core SNP-based phylogenetic tree clade designations (based on Figure 3). Colored stars indicate country of origin for each human-source isolate (USA-blue, Canada-red). The outer four rings denote the presence of genetic elements *sopE*, *uidA*, full *cirA*, frameshifted *cirA*, and truncated *cirA*. The tree is rooted with a turkey-source isolate collected in 2002 (SRR1195634).

**Figure S6.** Schematic depicting quality filtering steps within the bioinformatic processing pipeline. Thirty-three isolates were removed during filtering for a final sample size of 988 high-quality *S*. Reading isolate genomes.

